# Bayesian Analysis of High Throughput Data

**DOI:** 10.1101/079525

**Authors:** Eric J. Ma, Islam T. M. Hussein, Vivian J. Zhong, Christopher Bandoro, Jonathan A. Runstadler

## Abstract

Duplicate or triplicate experimental replicates are commonplace in the high throughput literature. However, it has not been tested whether this is statistically defensible or not. To address this issue, we use probabilistic programming to develop a simple hierarchical model for analyzing high throughput measurement data. With the model and simulated data, we show that a small increase in replicate experiments can quantitatively improve accuracy in measurement. We also provide posterior densities for statistical parameters used in the evaluation of HT data. Finally, we provide an extensible open source implementation that ingests data structured in a simple format and produces posterior densities of estimated measurement and assay evaluation parameters.

## Introduction

High throughput (HT) screening experiments are necessary for systematically interrogating biology. However, there are a number of statistical issues that are widespread in the HT screening literature. Firstly, triplicate (or worse, duplicate) measurements are commonplace, with little statistical justification; conceivably, this is mostly cost driven, a practical reason but nonetheless detrimental for scientific accuracy. Secondly, t-tests with multiple hypothesis correction serves as the main vehicle for statistical analysis of HT data, potentially falsely identifying samples as negatives or non-hits. Thirdly, standard error of the mean (SEM) are commonly used as the reported error bars, not only in the HT literature but also in non-HT publications (Kemnitzer et al. 2005; Marion et al. 2009; Le Hellard et al. 2002; Fu et al. 2014), and this under-represents the variation in the data. Finally, statistical parameters for assay evaluation are computed without acknowledging the uncertainty that may arise because of uncertainty in the data.

To address these problems, take an empirical approach. We use probabilistic programming to develop a simple Bayesian hierarchical model of a ‘generic’ HT assay (Figure 1, Supplementary Materials). Using this model, we are able to simultaneously provide Bayesian posterior distributions of measurement and statistical evaluation parameters. We show, using simulation studies, that increasing the number of replicates by one or two measurements can drastically reduce measurement inaccuracy. Using both simulation and real data, we show that the common practice of reporting mean ± SEM under-represents the uncertainty in measurement variation. We argue that the uncertainty in statistical assessment parameters can help guide more rational decision-making. Finally, we provide an extensible open source tool for the analysis of such data.

**Figure 1:**
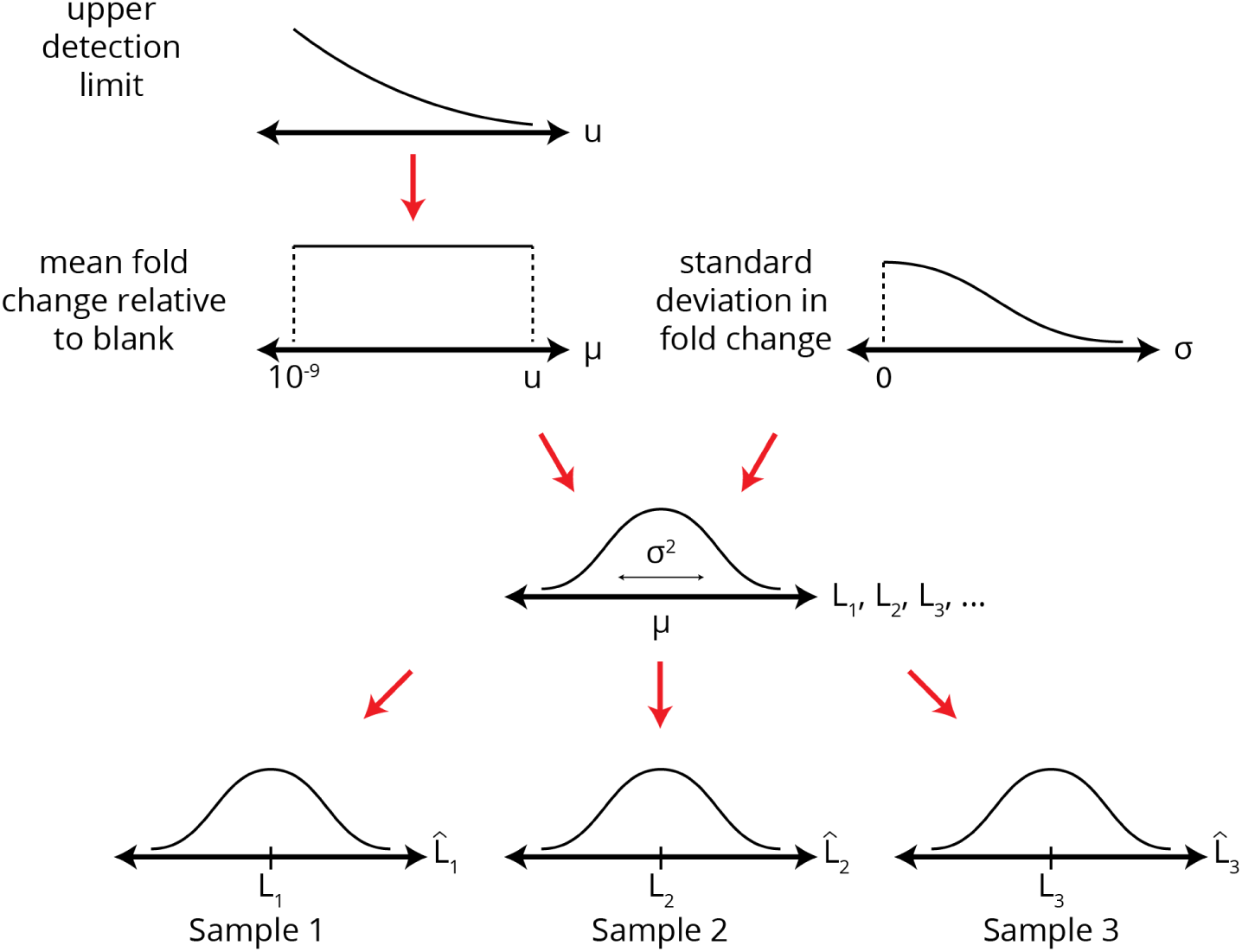
Bayesian hierarchical model.

## Results

### Statistically Defensible Replicate Measurements

In order to investigate how the number of replicates affected the accuracy, we simulated experimental runs of 100 samples with varying numbers replicate measurements (`n=2` to `n=20`). For each n, 20 experimental runs were simulated.

As shown in Figure 2, the baseline accuracy rate with duplicate (`n=2`) measurements, as measured by fraction of actual values inside the posterior density’s 95% HPD, falls around the 70-75% range. This means that about 25% of the final posterior 95% HPDs do not encompass the actual value. By contrast, by using `n=5` replicates, the accurate HPD fraction falls around the 85-90% range. Roughly doubling the number of samples decreases the inaccurate fraction by up to 3-fold. Following the law of diminishing marginal returns, additional accuracy can be gained, but at a cost of increasing sample sizes.

**Figure 2:**
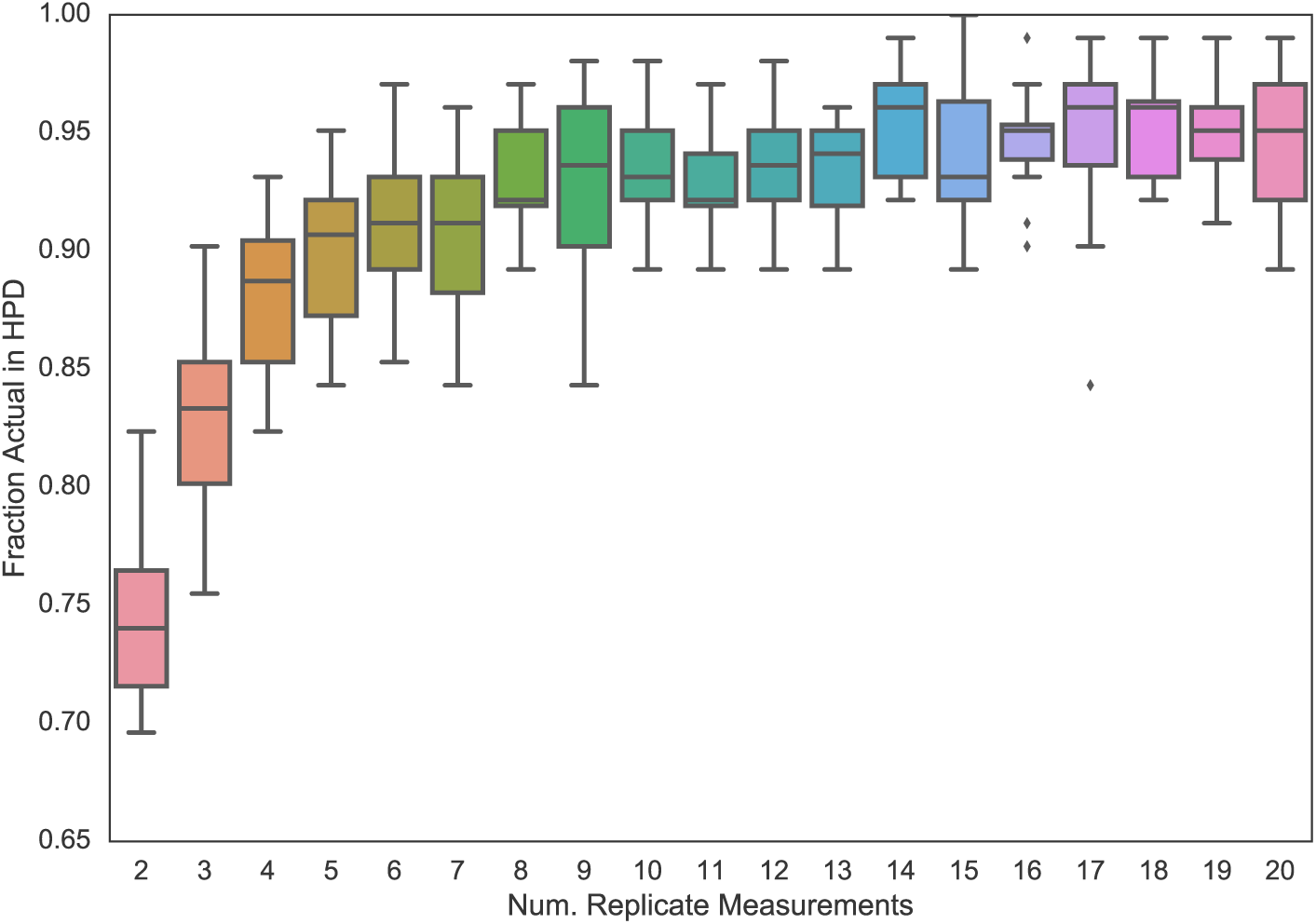
Accuracy of 95% HPD as a function of number of replicate samples taken.

## Representation of Uncertainty

Statistical software (e.g. GraphPad Prism) make it easy for researchers to visualize and compute frequentist confidence intervals and error bars. However, as a result of their ease of use, it is also easy to make statistical errors such as reporting error bars using the standard error of the mean (SEM), rather than 95% confidence/credible intervals. Our analysis of simulation and experimental data show clearly what can be inferred from the mathematical form but is often ignored: that the SEM grossly under-represents the uncertainty in measurement and data variation compared to 95% confidence intervals and Bayesian 95% credible intervals (Figure 3). As such, it would be poor statistical practice to report SEM, and 95% credible/confidence intervals would be much preferable.

**Figure 3:**
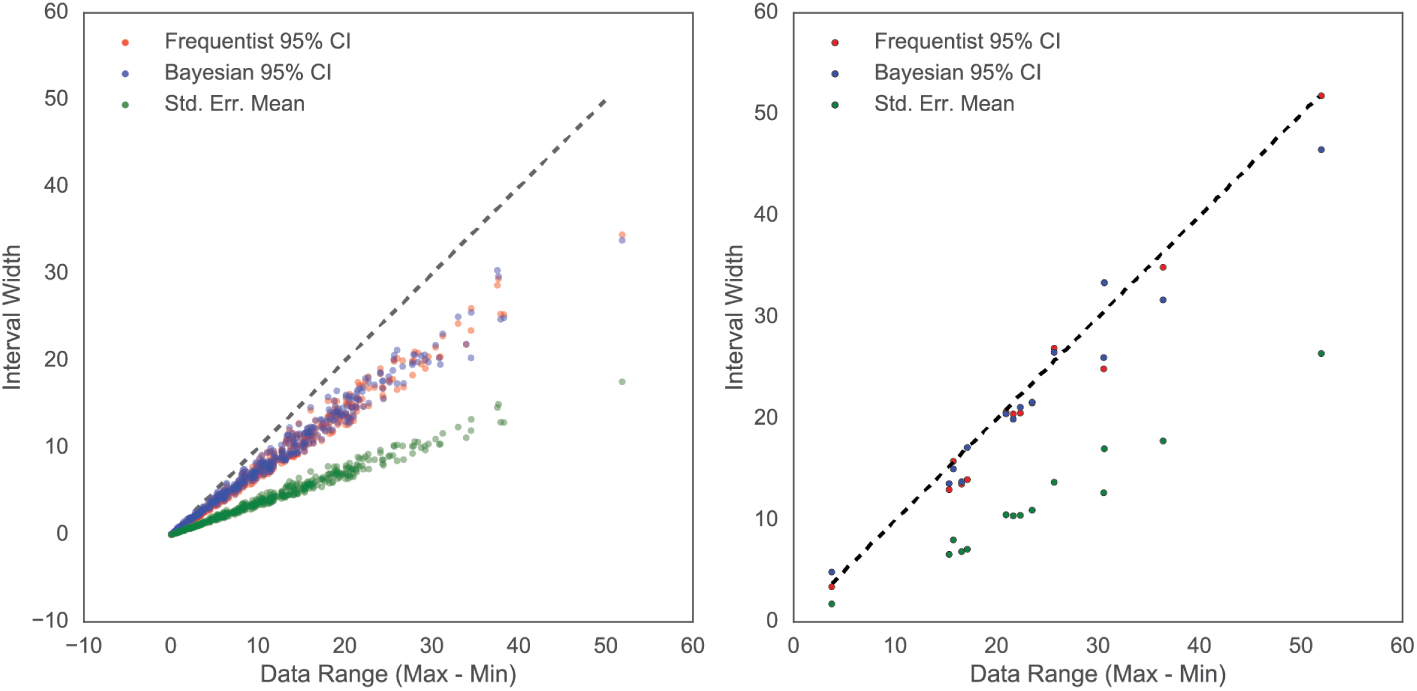
Bayesian 95% credible interval, Frequentist 95% confidence interval, and SEM interval widths as compared to the actual data range. (left) Simulated data. (right) Experimental measurement data measuring the influence of heat-killed bacteria on influenza activity.

## Posterior Densities of Assay Parameters

Statistical parameters, such as the Z-factor, have been developed to evaluate the quality of HT assay data (J. Zhang, Chung, and Oldenburg 1999; Sui and Wu 2007). By taking a Bayesian view, we can compute not just the expected parameter values but also their posterior distributions (Figure 4). As such, given the uncertainty surrounding the measurements, the original 3-class system for classifying the quality of an hit can be extended to 5 classes (Figure 4).

**Figure 4:**
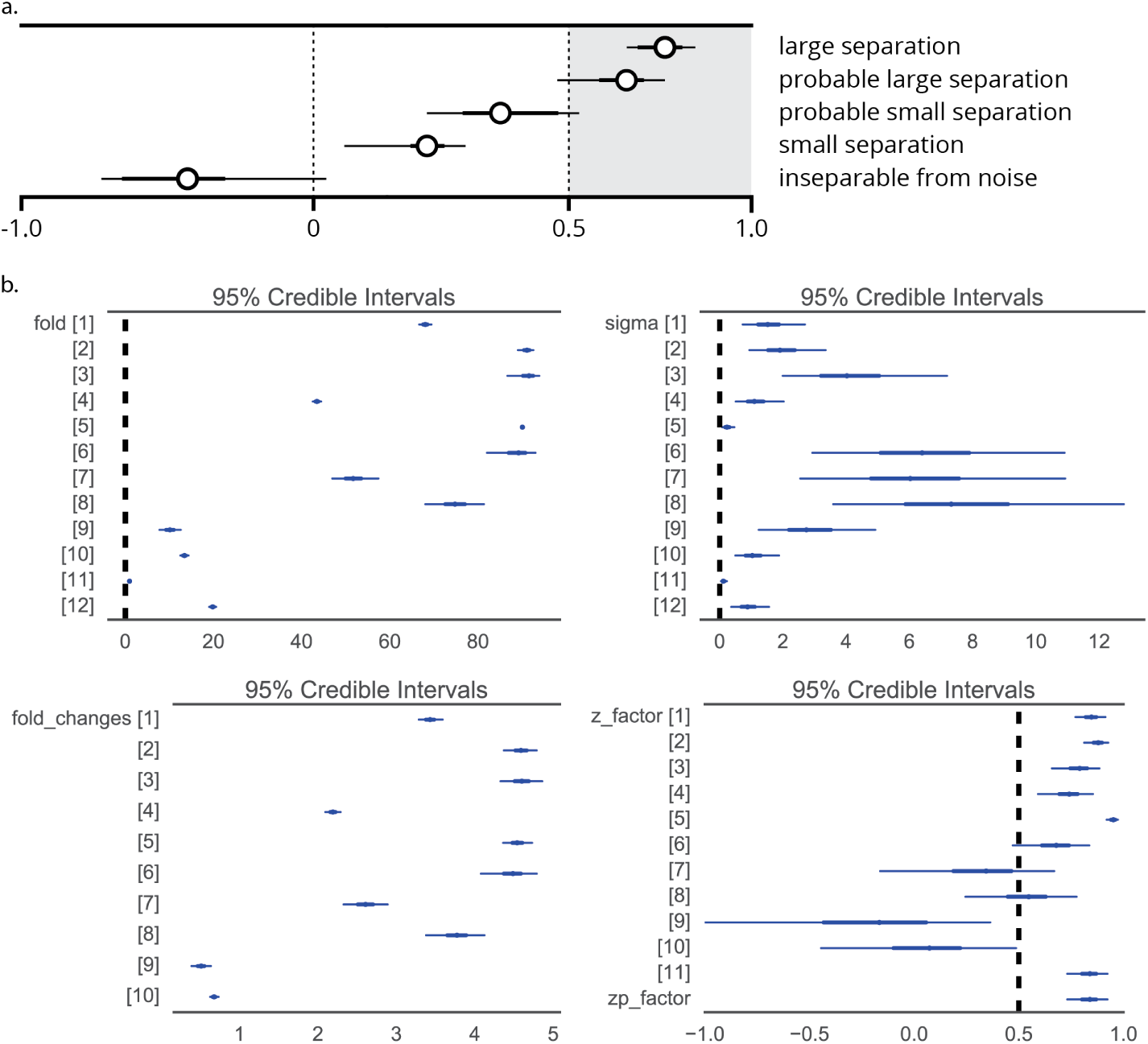
Z-score classes and simulation data. Circle/dot: HPD mean. Thick lines: HPD inter-quartile range. Thin lines: 95% HPD range. (a) Five Z-score classes based on the Z-score posterior density. (b) Forest plot of posterior distributions from one simulation run. Samples 11 and 12 (respectively) are the blank and the non-extreme positive control in this simulated experiment. (top-left) Posterior density in fold change relative to blank. (top-right) Posterior density of variance. (bottom-left) Deterministic posterior density of fold change relative to positive control. (bottom-right) Deterministic posterior density of Z-factor computed using the non-extreme positive control as the baseline.

The actionable consequences of these Z-value distributions depends on the experimental context. There may be scenarios where downstream experimentation is expensive, and only “true hits” should be tested; in this case, the “probable large separation” samples may be chosen for exclusion, helping to reduce costs. On the other hand, if downstream experimentation is cheap, and it is desirable to have a large set of samples to be processed further, then samples in the “probable small separation” may be included in downstream testing, helping to reduce false negatives. The truism remains: statistics does not replace human judgment of the value of a sample, but can serve as a valuable tool in the decision-making process.

We note that Z-factors are not the only statistical parameters that can be computed. Other deterministically calculated parameters, such as effect sizes, can be computed in a similar fashion, likewise yielding uncertainty estimates, given the data.

## Discussion

It is well-known that Bayesian analysis allows the uncertainty in parameter estimates to be explicitly modelled, with credibility (probability density) assigned to parameter estimate intervals. The provision of uncertainty can clarify close-to-call situations (e.g. Z-factors close to 0.5) and uncover potential false positives (e.g. large Z-factors close to 1.0 but with high variance), enabling better decision-making under uncertainty. Other merits and caveats of Bayesian analysis have been treated extensively in the literature as well (Kruschke 2013; Lin et al. 1999), and we do not go further into them here.

Probabilistic programming approaches make Bayesian methods much more accessible than analytical methods (Salvatier, Wiecki, and Fonnesbeck 2015). By leveraging these tools, we are in turn able to make Bayesian methods more generally accessible for the generic researcher working in high throughput measurement. In aid of reproducible science, we have also released an open source command-line program available for analysis of this type of data (#cite: Zenodo).

A key issue that has cropped up over the past half decade is the scientific “reproducibility crisis”, partly due to erroneous researcher reliance on p-values as a judgement device for “significance” (Wasserstein and Lazar 2016). Judgements of what “hits” to continue with downstream processing often relies on a calculated p-value rather than effect sizes; statistical significance has come to replace biological significance (Nuzzo 2014; Baker 2016). In light of this, we argue that by taking a Bayesian view of the data, we may replace p-value-based judgement calls with ones based on the distribution and uncertainty in estimated quality evaluation parameters (e.g. Z-factors & effect sizes), hence improving the quality of published results in the scientific literature.

## Materials and Methods

### Code & Data

All code for simulation and analysis are available as Python scripts and Jupyter notebooks. The archived version used in this publication is released on Zenodo (#TODO), while the source code and data (including that used for this manuscript) can be found on GitHub.

